# MUSICiAn: Genome-wide Identification of Genes Involved in DNA Repair via Control-Free Mutational Spectra Analysis

**DOI:** 10.1101/2025.01.27.635038

**Authors:** Colm Seale, Marco Barazas, Robin van Schendel, Marcel Tijsterman, Joana P. Gonçalves

## Abstract

**Motivation:** Understanding the factors involved in DNA double-strand break (DSB) repair is crucial for the development of targeted anti-cancer therapies, yet the roles of many genes remain unclear. Recent studies show that perturbations of certain genes can alter the distribution of sequence-specific mutations left behind after DSB repair. This suggests that genome-wide screening could reveal novel DSB repair factors by identifying genes whose perturbation causes the mutational distribution spectra observed at a given DSB site to deviate significantly from the wild-type. However, designing proper controls for a genome-wide perturbation screen could be challenging. We explore the idea that a genome-wide screen might allow us to forgo the use of traditional non-targeting controls by reframing the analysis as an outlier detection problem, assuming that most genes have minimal influence on DSB repair.

**Results:** We propose MUSICiAn (Mutational Signature Catalogue Analysis), a compositional data analysis method that ranks gene perturbation-specific mutational spectra without controls by measuring deviations from the central tendency in the distributions of all spectra. We show that MUSICiAn can effectively estimate pseudo-controls for the existing Repair-seq dataset, screening 476 genes and 60 non-targeting controls. We further apply MUSICiAn to a genome-wide dataset profiling mutational outcomes induced by CRISPR-Cas9 at three target sites across cells with individual perturbations of 18,406 genes. MUSICiAn successfully recovers known genes, highlights the spliceosome as a lesser-appreciated player in DSB repair, and reveals candidates for further investigation.

**Availability:** github.com/joanagoncalveslab/MUSICiAn.

## Introduction

Double-strand breaks (DSBs) in DNA are critical cellular events that occur spontaneously due to endogenous processes like replication or external agents like ionizing radiation. Left unaddressed, DSBs can lead to genomic instability and eventually cell death or cancer (Ceccaldi *et al*., 2016). As a result, cells have evolved a suite of mechanisms to repair DSBs, including the non-homologous end joining (NHEJ), microhomology-mediated end joining (MMEJ, also called alt-NHEJ), and homology-directed repair (HDR) pathways (Scully *et al*., 2019). Understanding the roles that genes play in DSB repair can importantly contribute to the development of targeted therapies for diseases such as cancer (Trenner and Sartori, 2019; Awwad *et al*., 2023). For example, PARP inhibitor drugs are indicated to treat cancers with impaired HDR or BRCA gene function, whose synthetic lethality with PARP is leveraged to block DSB repair and cause a fatal accumulation of DNA damage in HDR- or BRCA-deficient cancer cells (Chen, 2011). The ability to discover further opportunities for targeted therapy requires deeper insight into gene function, yet for many genes the link with DSB repair remains unclear.

In searching for these links, research has turned to large-scale gene functional screens enabled by CRISPR technology (Awwad *et al*., 2023). Originally, functional screens for DSB repair focused on the effect of gene silencing or knockout on readouts such as cell growth and proliferation to identify repair factors (Hurov *et al*., 2010; Smogorzewska et al., 2010; Zimmermann et al., 2018; O’Connell *et al*., 2010; Olivieri *et al*., 2020). While valuable to characterize gene essentiality, inhibition of cellular growth is only indirectly related to DSB repair and could lead to results confounded by other mechanisms of cellular activity. For more precise readouts and biological insights, recent advances use CRISPR targeting to induce DSBs and deep sequencing to analyze how disruption of gene function alters the mutational spectra, or the frequency distributions of mutations arising at DSB sites following repair (van Overbeek *et al*., 2016; Wyatt *et al*., 2016; Bothmer et al., 2017; Shen et al., 2018; Shou et al., 2018; Hussmann et al., 2021). Multiple studies have demonstrated that knockouts of certain genes yield distinct, sequence-specific mutational spectra (van Overbeek *et al*., 2016; Shen *et al*., 2018; Hussmann *et al*., 2021), but focused on the screening of known DSB repair genes. Notably, the first genome-wide study characterizing the effect of gene perturbation on mutational spectra will soon be released. We obtained early access to the data from this study, termed Mutational Signature Catalogue (MUSIC), to be made available upon publication.

Using CRISPR targeting with mutational spectra as readout, the primary approach to link genes to DSB repair is to quantify how much the mutational spectrum deviates from the expected wild-type distribution following the knockout of each individual gene. The larger the deviation, the higher the confidence that the perturbed gene has an effect on the outcomes and could thus be involved in DSB repair. Recent work by Hussmann *et al*. (2021) defined this deviation as the “overall outcome redistribution activity”, quantified by a chi-squared-like statistic relying on non-targeting controls to determine the expected wild-type spectrum. For genome-wide screens, a limited set of non-targeting controls might not be suitable. While the majority of genes is not expected to produce an effect on the mutational spectra, it is unclear if targeting such genes could indirectly or mildly influence the outcomes, an effect which would not be appropriately captured by non-targeting controls. At the same time, it would be challenging to design realistic controls for all levels of variation at play in a genome-wide screen, while trying to maximize the coverage per mutational spectra and mitigate batch effects. We explore an alternative approach leveraging the assumption that most genes in a genome-wide perturbation screen have minimal impact on the mutational spectra to frame the identification of DSB repair genes as an outlier detection problem (Aggarwal, 2016), and investigate if it can forgo the need for conventional controls.

When analyzing mutational spectra, it is also important to consider their compositional nature. In other words, each mutational spectrum is a distribution of relative frequencies over a collection of mutation categories whose overall sum is one. This composition property introduces a negative correlation bias caused by dependencies between the different frequencies, where an increase for one mutation type necessarily causes a reduction in others. Ignoring the dependencies in compositional data using standard data analysis techniques can lead to misleading results and interpretation (Aitchison, 1983). Additionally, the covariance structure of mutational spectra is likely to be skewed by the outlier gene knockouts that significantly affect DSB repair, emphasizing the need for methods tailored for compositional data analysis.

We introduce MUSICiAn (<u>M</u>utational <u>S</u>ignature <u>C</u>atalogue <u>A</u>nalysis), a computational approach to score gene associations with DSB repair via genome-wide mutational spectra analysis. MUSICiAn operates without non-targeting controls, framing the task as an outlier detection problem under the assumption that most genes do not influence DSB repair. MUSICiAn uses the compositional data analysis (CoDA) framework to address dependencies and outliers in genome-wide mutational spectra data, for an improved estimation of pseudo-controls. By ranking gene knockouts based on their robust deviation from the overall mutational spectra distribution, MUSICiAn provides a control-free approach for genome-wide discovery of DSB repair-related genes.

We evaluate the MUSICiAn estimation of pseudo-controls on the Repair-seq dataset, screening 476 DSB genes and 60 non-targeting controls (Hussmann *et al*., 2021). We further apply MUSICiAn to the genome-wide MUSIC mutational spectra dataset, covering 18,406 genes, to investigate the ability of this control-free method to recover established repair genes and suggest new candidates for experimental validation.

## Methods

We introduce the MUSICiAn method using outlier detection to identify DSB repair genes from genome-wide CRISPR mutational spectra without traditional controls. The aim is to quantify the effect that each gene knockout produces on the mutational spectra relative to the expected wild-type or control spectra. In the absence of controls, MUSICiAn leverages the assumption that most genes are not involved in DSB repair to estimate the center of the mutational spectra distribution as a representative point, close to which the spectra will be most alike the expected wild-type. To quantify the deviation, MUSICiAn calculates a distance between each spectra and the estimated center also taking the covariance of the spectra distribution into account. This is done using a combination of data transformation and robust covariance estimation designed to address dependencies and outliers in the mutational spectra data. Finally, MUSICiAn creates a unified gene outlier score based on the distances obtained across target sites.

### Data and preprocessing

#### Mutational outcome data

We analyzed data from two gene perturbation screens with CRISPR induced mutational outcome readout, Repair-seq and MUSIC (Supplementary Fig. S1 for an illustration of the experimental setup). The Repair-seq screen used CRISPR interference with each of 1,573 single-guide RNAs (sgRNAs) to individually silence each of 476 DSB repair genes, and 60 non-targeting control sgRNAs (Hussmann *et al*., 2021). To generate mutational outcomes, Repair-seq used CRISPR-Cas9 to create DSBs for a single target site across the population of cells with and without silenced genes, in two biological replicates. The genome-wide MUSIC screen was similarly set up, but used CRISPR knockouts rather than interference, with 89,571 sgRNAs spanning 18,406 genes, and generated outcomes for three target sites in two biological replicates each. We downloaded the raw Repair-seq sequence data (Hussmann *et al*., 2021) from the NCBI Sequence Read Archive, Bioproject PRJNA746980, runs SRR15164738 and SRR15164739. We also obtained early access to the MUSIC sequence data, to be made available upon publication.

We called mutations from the sequence data using the Sequence Interrogation and Quantification (SIQ) tool (van Schendel *et al*., 2022) v4.3 with parameters “-m 2 -c -e 0.05”, specifying a minimum number of 2 reads for counting an event, the collapsing of identical events to a single record with the sum of counts, and a maximum permitted base error rate of 0.05. The SIQ tool mapped the reads to the sgRNAs used for gene perturbation and identified mutations observed at the CRISPR-Cas9-induced DSB sites.

#### Mutation aggregation and categorization

To reduce sparsity and improve statistical power, the fine-grained mutational outcomes output by the SIQ tool were aggregated into 8 higher-level categories: *wild-type*, denoting a sequence without mutations; *deletion with 1\2\3+bp\no microhomology*, for a deletion overlapping the cut site with a microhomology (MH) of length 1bp to 3+bp or no MH at all, where an MH is a short homologous sequence on both sides of the DSB and used for repair by MMEJ (Lieber, 2008); *insertion*, for a new sequence added at the cut site; *deletion with insertion*, for a combination of deletion and insertion; and *homology-directed repair*, for any insertion matching the donor template DNA (Supplementary Table S3 for SIQ vs. MUSICiAn categories). Any other rare mutation types, such as single-base substitutions, were excluded, since they are not typical outcomes of DSB repair. Wild-type reads were also excluded to avoid confounding by gene essentiality, as a decrease in wild-type read abundance could indicate that the gene was essential for survival, but not if it was relevant for DSB repair. We thus considered a final set of 7 mutation categories.

#### Quality analysis and filtering of perturbation sgRNAs

For each replicate, we filtered out perturbation sgRNAs yielding a total read count below 700 across the 7 mutation categories, and excluded genes with less than 3 associated perturbation sgRNAs. Additionally, we controlled for inconsistencies in the effect of the different sgRNAs used for perturbation of the same gene, which could be indicative of sgRNA off-target effects, less effective gene perturbation, or any other undesirable effect. We excluded sgRNAs whose count profiles over the 7 mutation categories showed a median pairwise Pearson’s correlation below 0.6 with the profiles for other sgRNAs perturbing the same gene within the same replicate (and target site), or a median pairwise Pearson’s correlation below 0.6 with replicate profiles for the same sgRNA and target site (Supplementary Tables S1 and S2). To avoid numerical issues with the data transformation applied by MUSICiAn later on, in the rare cases where some mutation categories had zero counts, we imputed real values drawn independently from a uniform distribution between the detection limit *DL* and 0.1 *× DL*, where *DL* = 1. (Lubbe *et al*., 2021).

#### Generating mutational spectra per gene

We first computed mutational spectra by dividing the count of each of the 7 mutation categories by the total per sgRNA and replicate. Then we aggregated across sgRNAs by calculating the geometric mean of the sgRNAs-associated spectra per gene and replicate, producing replicate spectra per gene (two for each target site). Finally, we computed the geometric mean between replicate spectra per target site, resulting in one mutational spectra per gene and target site. After every aggregation step, the frequencies in each mutational spectra were divided by the sum to make sure they summed to one.

The MUSICiAn method scores genes for DSB repair association by computing the distance between the mutational spectrum of each perturbed gene and the estimated center of the distribution of all spectra obtained for a given target site (Fig. 1). For experiments with multiple targets, target-specific scores can be normalized and averaged to produce one single gene score. Genes with larger scores have a more prominent effect on the mutational spectra, thus also a higher likelihood of being involved in DSB repair. Genes with the lowest scores are assumed to approximate the wild-type or control distribution. We are interested in both the most outlying and the most central spectra for downstream analysis.

**Fig. 1:**
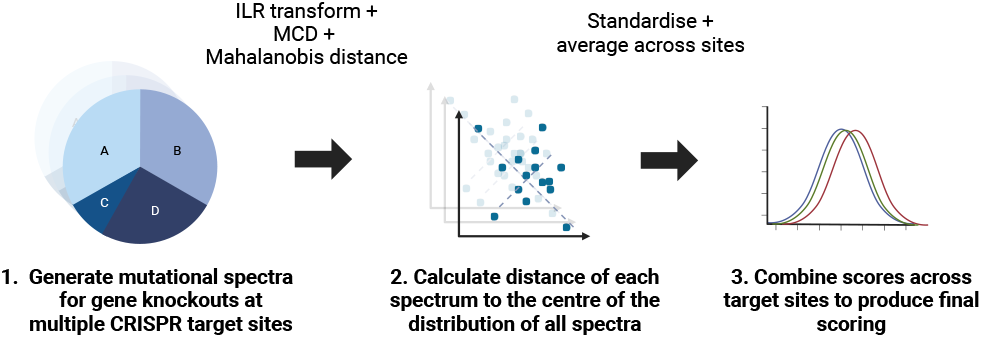
Overview of MUSICiAn scoring of gene effect on mutational spectra. The method quantifies the effect of gene perturbation using the Mahalanobis distance of the gene mutational spectrum to the estimated center of the spectra distribution of all genes, under the assumption that most genes have a negligible effect. Estimation is improved using ILR-transformed spectra and robust covariance (MCD) to mitigate biases from data closure and outliers. Distances are normalized and averaged across target sites to produce a unified gene effect score.

#### Gene scoring

To calculate gene effect scores, MUSICiAn computes the Mahalanobis distance (Mahalanobis, 2018) per gene spectrum relative to the overall spectra distribution per target site, assuming that most genes are not directly involved in DSB repair and have negligible effect (Fig. 1). We chose the Mahalanobis distance as it takes the distribution and covariation of the data into account, unlike the Euclidean distance. Informative Mahalanobis distances require reliable covariance estimation, which is affected by: data closure, where dependencies between spectra categories summing to one introduces a negative correlation bias (Aitchison, 1983); and outlier genes with a significant impact on mutational spectra and therefore also on the distribution. To mitigate data closure, MUSICiAn applies an isometric log-ratio (ILR) transformation (Egozcue *et al*., 2003) to the mutational spectra using the defaults for scikit-bio 0.5.4. The ILR transformation maps the data from a constrained simplex space to an unconstrained Euclidean space, allowing for independent statistical analysis of components. To mitigate outlier effects, MUSICiAn uses the minimum covariance determinant (MCD) as a robust covariance matrix estimator (Filzmoser *et al*., 2009), using scikit-learn 1.2.1 defaults.

The MUSICiAn method calculates the robust Mahalanobis distances for each ILR-transformed spectra, and unifies the individual distances into gene scores across target sites by: *(i)* selecting the common genes with mutational spectra in all target sites to act as a reference, *(ii)* calculating the mean and standard deviation of the distances of the reference genes per target site, *(iii)* normalizing all gene distances per site by subtracting the means and dividing standard deviations to place them on a common scale, and *(iv)* averaging the normalized gene distances across sites, ignoring missing values, to produce a final unified gene score.

#### Pseudo-control selection

The target-specific distances and unified gene scores enable the selection of “pseudo-controls” as the lowest-scoring genes per target or common across target sites. These pseudo-controls enable comparative analyses by estimating the central tendency of traditional controls, but may not recapitulate their natural variation.

### Evaluation

Before applying MUSICiAn to the genome-wide MUSIC dataset, we assessed the outlier detection and pseudo-control selection on the Repair-seq dataset, the only other dataset available of CRISPR mutational spectra for multiple individual gene knockouts. While not a genome-wide dataset, Repair-seq included non-targeting controls, allowing us to assess if and how well MUSICiAn could estimate the wild-type distribution center. Furthermore, Repair-seq data focused on genes involved in DNA repair, so we also checked if MUSICiAn could recover similar mutational patterns for the genes screened in both studies. We preprocessed the Repair-seq data as described and held out the non-targeting controls from the scoring for later validation.

#### Estimation of pseudo-controls

We used PCA to visualize the effect of ILR transformation and robust MCD covariance, proposed to mitigate compositional data closure and outlier spectra, on the estimation of the mutational spectra distribution center and selection of pseudo-controls for the Repair-seq dataset. We applied PCA in four scenarios: *Classical Covariance*, using the original mutational spectra with the classical covariance estimation; *MCD Covariance*, using the original spectra with the outlier-robust MCD covariance estimation; *ILR & Classical Covariance*, using ILR-transformed spectra with classical covariance estimation; and *ILR & MCD Covariance*, using ILR-transformed spectra with MCD covariance estimation. After ILR transformation, location and covariance estimation, we back-transformed the data to centered log-ratio (CLR) space to analyze the relation between PCA components and mutation categories (Filzmoser *et al*., 2009). To evaluate pseudo-control selection, we identified 60 pseudo-controls for each scenario and calculated the Jensen-Shannon distance (JSD) between the geometric means of the non-targeting control and the pseudo-control spectra. As a baseline, we also calculated the distance from the non-targeting control spectra to the geometric mean across all gene-targeting sgRNAs, without pseudo-control selection. The JSD quantifies the distance between distributions, where a lower distance indicates greater similarity between distributions.

#### Cross-dataset estimation of pseudo-controls

To further assess the selection of pseudo-controls, we analyzed the consistency in mutational patterns retrieved for the MUSIC and Repair-seq datasets, using pseudo-controls estimated by MUSICiAn jointly from the two datasets. Specifically, we applied MUSICiAn to select 60 pseudo-controls for the set of all mutational spectra associated with the 434 genes shared across both datasets, with four target sites in total (three for MUSIC, one for Repair-seq). We then calculated the difference in mutation frequency per category between each gene-related mutational spectra and the geometric mean of the pseudo-controls. Finally, we performed hierarchical clustering (Ward Jr, 1963) on the resulting difference matrix, using Ward cluster linkage and distance between samples based on Pearson’s correlation.

#### Gene scoring and ranking performance

To evaluate the quality of the MUSICiAn-derived gene effect scores for the genome-wide MUSIC dataset, we examined if MUSICiAn could effectively recover genes with known links to DSB repair by scoring or ranking them higher than other genes based on their effect on the mutational spectra. We assessed performance separately using precision-recall (PR) curves against known DSB repair genes from two sources: 476 experimentally validated genes curated by Repair-seq for their AX227 CRISPRi library (Hussmann *et al*., 2021), and 295 genes whose annotations matched the regex “double-strand break repair|interstrand cross-link repair” (interstrand cross-link repair genes often crosstalk with DSB repair pathways such as HR, Michl *et al*. (2016)) in any field in the Gene Ontology (Ashburner *et al*., 2000; Consortium, 2021). For baseline comparison, we calculated PR curves after randomly ranking all genes in the MUSIC dataset. We preferred PR rather than ROC curves, given that the dataset is highly imbalanced, where most genes have no known association with DSB repair and are therefore considered negative for the purpose of the evaluation.

#### Functional enrichment for top 500 ranked genes

We performed enrichment analysis for the top 500 genes ranked by MUSICiAn against the background of all genes in the MUSIC dataset, using the “gseapy” python package 1.0.4. We employed four sets of annotations, including KEGG pathways “KEGG 2019 Mouse” (Kanehisa *et al*., 2023), and Gene Ontology terms across the three ontologies “GO Biological Process 2023”, “GO Molecular Function 2023”, “GO Cellular Component 2023”. We performed a hypergeometric test per term within each gene set, and the resulting *p*-values were FDR corrected using the Benjamini-Hochberg method. (Benjamini and Hochberg, 1995). We further estimated the effect of the genes annotated with each of the top 10 enriched terms or pathways on the mutation frequencies separately for the 4 annotation sets. To do this, we fitted an ordinary least squares (OLS) regression model per term *t* and mutation outcome category *o* to explain the variation in mutation frequency (*Frequency*) based on term or pathway membership (*Group*), according to the following R-style formula

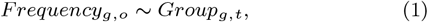

where each sample is a mutational spectra for a given gene knockout *g*. The *Frequency*_*g*,*o*_ variable denotes the frequency of the given mutation outcome *o* for gene *g*, and *Group*_*g*,*t*_ is a binary variable indicating if gene *g* is a member of term or pathway *t*. As case samples, we took the mutational spectra of all genes annotated with the enriched term in question. As control samples, we used the set of 100 pseudo-controls or lowest scoring genes, with valid mutational spectra across all target sites, and that were not members of any of the enriched pathways. We used the same control samples for the regression analysis of every annotation set, and report the regression coefficients and Benjamini-Hochberg corrected p-values for the *Group* variable.

## Results

### MUSICiAn can estimate absent control mutational spectra

We first assessed the ability of MUSICiAn to estimate pseudo-control mutational spectra in the absence of actual controls. To do this, we applied MUSICiAn to the Repair-seq dataset, containing mutational spectra for one target site across knockouts of 476 different genes and 60 actual non-targeting controls. The actual controls were left out to be able to quantify how well they could be recovered by MUSICiAn. We also isolated the contributions of the ILR transformation and robust covariance (MCD) used by MUSICiAn to investigate if they improved the estimation of the distribution center location and covariance, and ultimately the selection of pseudo-controls, in the presence of outlier spectra and negative correlation bias.

To visually examine the effect of ILR transformation and MCD on the distribution, we applied PCA to the original and ILR-transformed mutational spectra separately using classical PCA and a robust variant of PCA based on the MCD. The estimated center of the distribution appeared to align the best with the center determined based on the actual controls (geometric mean of the 60 non-targeting controls) when both ILR and MCD were used to respectively address data closure and outliers in the mutational spectra data (Fig. 2A, “ILR & MCD Covariance” vs. others). We further observed that the pseudo-controls selected as the 60 mutational spectra closest to the center of the distribution estimated by MUSICiAn, using any of the four combinations of spectra and covariance types, were far more similar to the actual non-targeting controls than the average across all spectra. Specifically, the Jensen-Shannon (JS) distances between the geometric means of pseudo-controls and non-targeting controls were one order of magnitude smaller than those between the geometric means of all spectra and non-targeting controls (respectively *<* 0.005 and 0.012, Fig. 2B). Moreover, the selection of pseudo-controls using the preferred combination of techniques in MUSICiAn, ILR transformation and robust MCD covariance, produced the closest match with the actual non-targeting controls than the other three (JS distances 3.73*×*10^−3^ against 3.77*×*10^−3^, 4.42 *×* 10^−3^, and 4.69 *×* 10^−3^; Fig. 2B). This result supported our choice to place ILR transformation and MCD at the core of the MUSICiAn outlier detection algorithm.

**Fig. 2:**
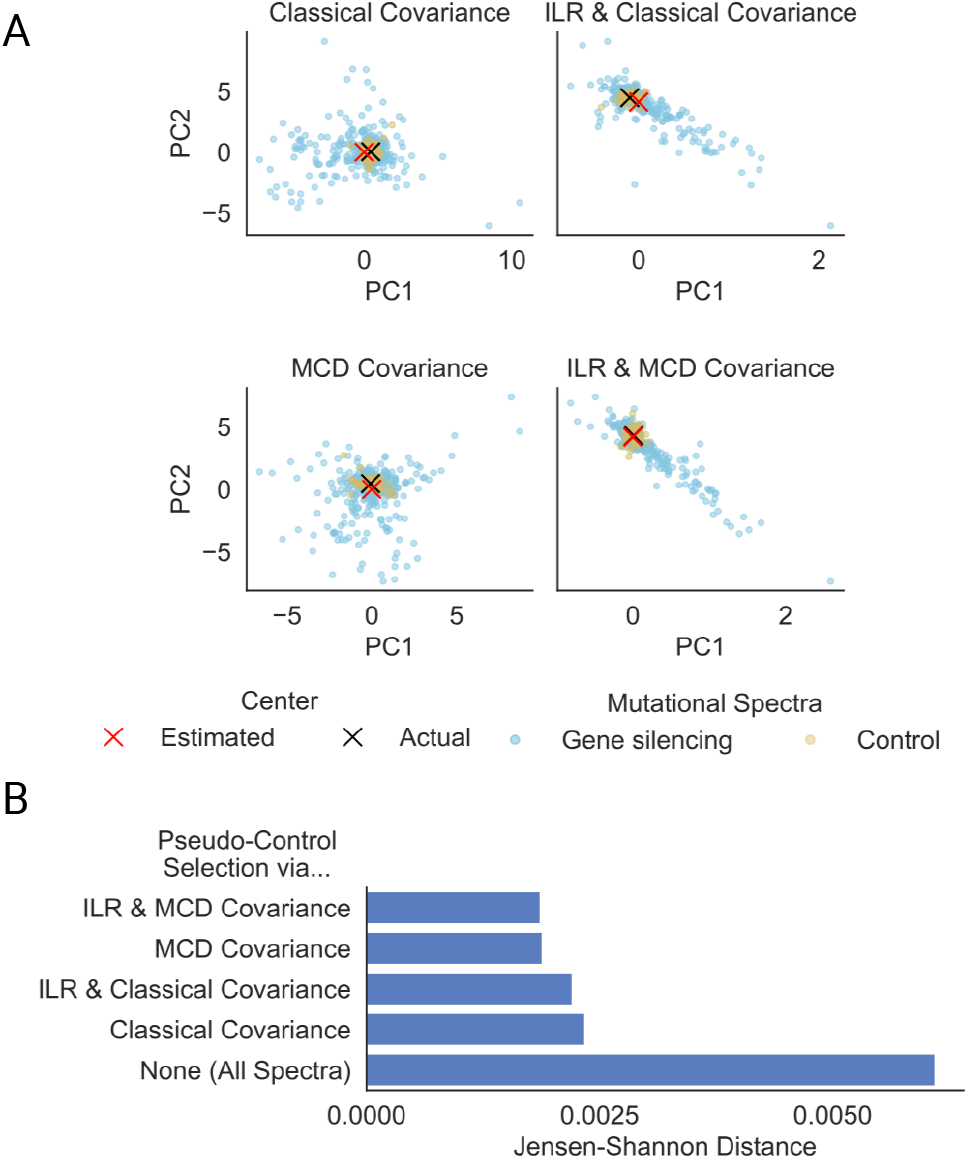
Evaluation of MUSICiAn-selected pseudo-controls based on the estimated mutational spectra distribution center for Repair-seq. Effect of ILR transformation and MCD covariance on (A) the estimated center of the mutational spectra distribution and (B) the selected pseudo-controls, using the original or ILR transformed spectra with classical or MCD covariance. For (A), actual center (black cross) of the mutational spectra distribution as the geometric mean of the 60 actual controls (yellow points), and center estimated by MUSICiAn (red cross) based on the mutational spectra under gene-silencing (blue points), projected onto the two axes of largest variation in the data (first two principal components). For (B), Jensen-Shannon distances between the geometric means of the 60 actual non-targeting controls and either all mutational spectra or the 60 pseudo-controls closest to the center estimated using each of the four combinations of spectra and covariance types.

We note that the majority of the 476 genes characterized in the Repair-seq screen are known to be involved in DNA repair, and therefore the assumption that most genes should not have an impact on the mutational spectra was in theory not necessarily met for this dataset. However, in practice, a large proportion of DNA repair genes still showed little effect on mutation frequencies (Fig. 2). The fact that MUSICiAn was able to recover controls in this scenario highlights that it could be applicable to more focused studies beyond genome-wide screens whenever a similar reasonable assumption can be made, for instance based on prior knowledge or the actual distribution of the data.

### MUSICiAn controls reveal known repair patterns across studies

We further questioned if MUSICiAn could estimate pseudo-controls for mutational spectra aggregated from different studies, such that consistent mutational repair patterns would be revealed when applying the same controls as a baseline across the studies. To address this, we jointly analyzed the mutational spectra for knockouts of the 434 genes screened in both the genome-wide MUSIC and the focused Repair-seq studies. After selecting pseudo-controls, we calculated the differences between the frequencies in each mutational spectrum, obtained under silencing or knockout of a specific gene, and the geometric mean of the pseudo-controls (Fig. 3). We also performed hierarchical clustering of genes and mutation categories based on those differences (Fig. 3). The results revealed consistency in how HDR and insertion events were influenced by silencing of specific genes across targets and studies, as well as broadly consistent patterns for other mutation types with larger variations that could be attributed to differing target site-specific characteristics within and between studies.

**Fig. 3:**
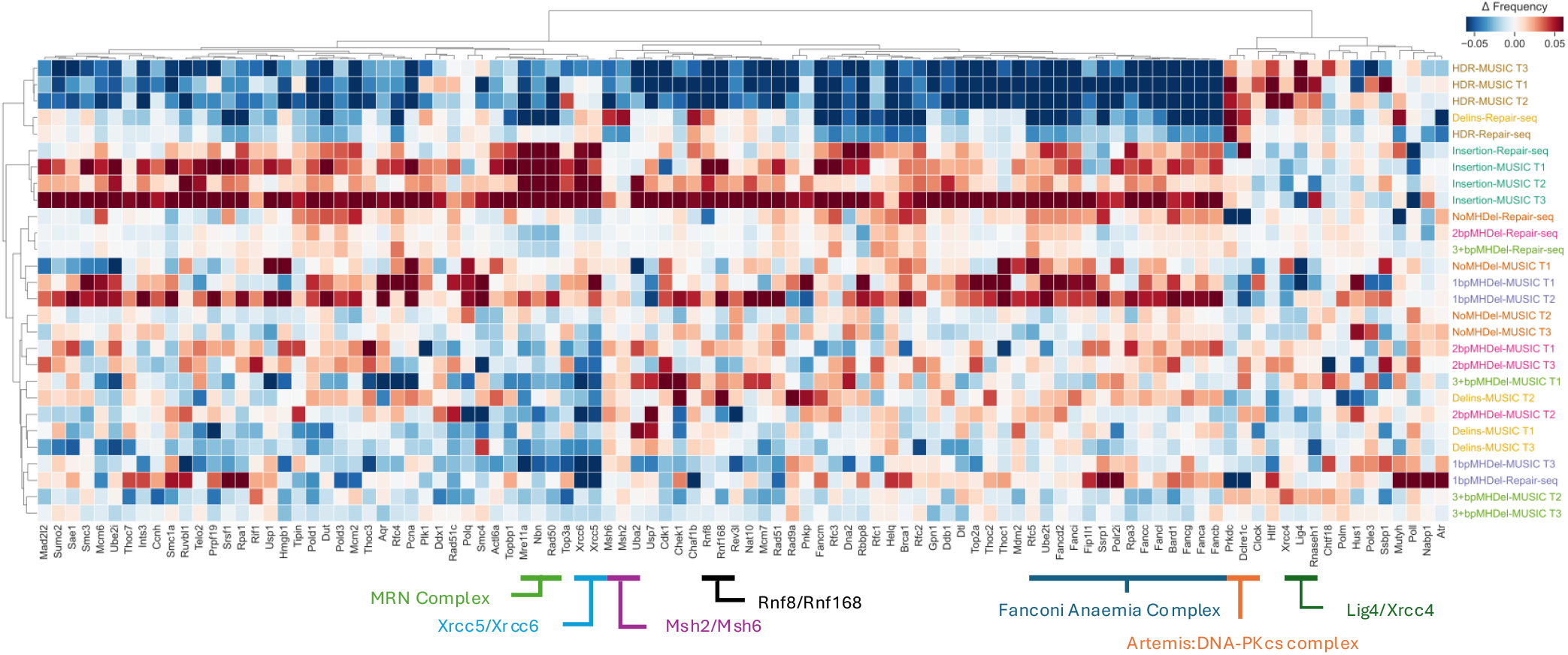
Heatmap of the difference in mutation frequencies between each spectrum obtained for the knockout of a specific gene and the geometric mean of the pseudo-control spectra selected by MUSICiAn, per mutation category and target site. Shown are the top 100 genes with the highest MUSICiAn outlier score across target sites (3 for MUSIC, denoted T1-T3, and 1 for Repair-seq). The horizontal axis represents genes. The vertical axis represents mutational outcomes, coloured by target site. Data was clustered on both dimensions, genes and mutation categories, using hierarchical clustering with Ward cluster linkage and distance between spectra based on Pearson’s correlation coefficient.

Gene clustering also identified meaningful groups, including the Fanconi anemia core complex and related genes, whose silencing suppressed HDR events (Fig. 3). Interestingly, *Helq* displayed a mutational pattern similar to these genes, suggesting a potential association with FA and HDR, a topic of ongoing debate (Jenkins *et al*., 2021; Thomas et al., 2022). Other notable clusters included: mismatch repair *MutS* homolog genes (*Msh2, Msh6*); ring finger protein genes with roles in DNA damage sensing and repair (*Rnf8, Rnf168*); NHEJ genes involved in early recognition of DNA damage and recruitment of additional repair factors (*Xrcc5, Xrcc6*), and in the processing of DNA ends (Artemis complex *Prkdc* and *Dclre1c*); and the MRN complex with roles in ATM checkpoint activation in response to DNA damage and also the tethering of broken DNA ends for further processing by NHEJ and HDR (*Mre11a, Rad50, Nbn*). The consistency in gene silencing effects on mutational spectra across the MUSIC and Repair-seq datasets, along with the identification of groups of genes with related function in DNA damage response, provided support for the effectiveness of the MUSICiAn control-free analysis in estimating pseudo-controls, quantifying effects, and ultimately generating meaningful insights from CRISPR targeting under gene silencing screens with mutational spectra readout.

### MUSICiAn recovers known gene-DSB repair associations

In addition to estimating pseudo-controls, MUSICiAn attributes an outlier score to each gene, which determines the multivariate effect of gene silencing on mutational spectra to suggest (novel) associations between the gene and DNA damage response. In this context, we first applied MUSICiAn to the genome-wide MUSIC dataset to assess if it could recover known repair genes. Genes were ranked by their MUSICiAn outlier score, and the ranking was evaluated against the set of 476 genes curated by Repair-seq and an alternative set of 295 genes retrieved from the Gene Ontology (GO). The closer to the top of the ranking these genes appeared, the better the results. We also performed the same evaluation on a randomly shuffled ranking as a baseline for comparison. The MUSICiAn method showed superior rankings for known associations with area under the precision-recall curve (AP) of 0.07 and 0.08 for the Repair-seq and GO gene sets, respectively, compared to an AP of 0.02 for the random baseline (Fig. 4).

**Fig. 4:**
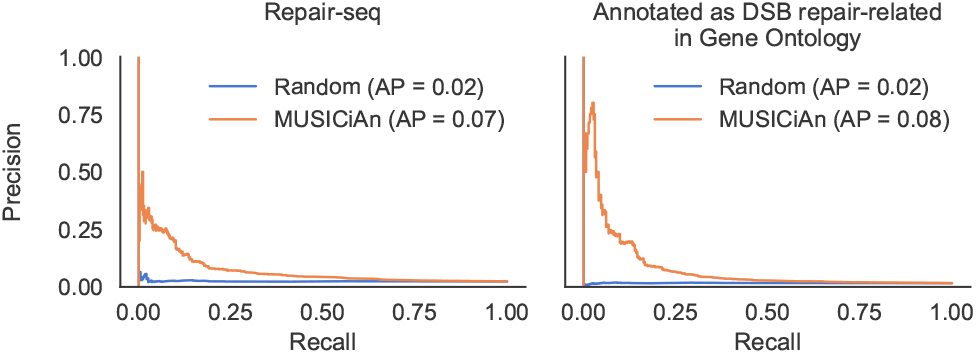
Performance of MUSICiAn, recovery of known DNA repair genes. Precision-recall curves using MUSICiAn ranking (orange) or random ranking (blue) against the following gold standards for evaluation: (left) curated genes in Repair-seq and (right) genes annotated with DSB repair-related in Gene Ontology.

Pathway enrichment analysis of the top 500 genes using KEGG annotations revealed significant associations with the “Fanconi anaemia” and “Homologous recombination” pathways (Fig. 5). A link with “Nucleotide excision repair” was also identified, supporting the idea that single and double-strand repair mechanisms are functionally intertwined (Zhang *et al*., 2009). Another enriched pathway, “Cell Cycle”, is known to influence DNA repair pathway choice (Zhang *et al*., 2009; Clay and Fox, 2021). Many DSB repair genes were also implicated in the “DNA replication” pathway (Burgers, 1998; Cortez, 2019).

**Fig. 5:**
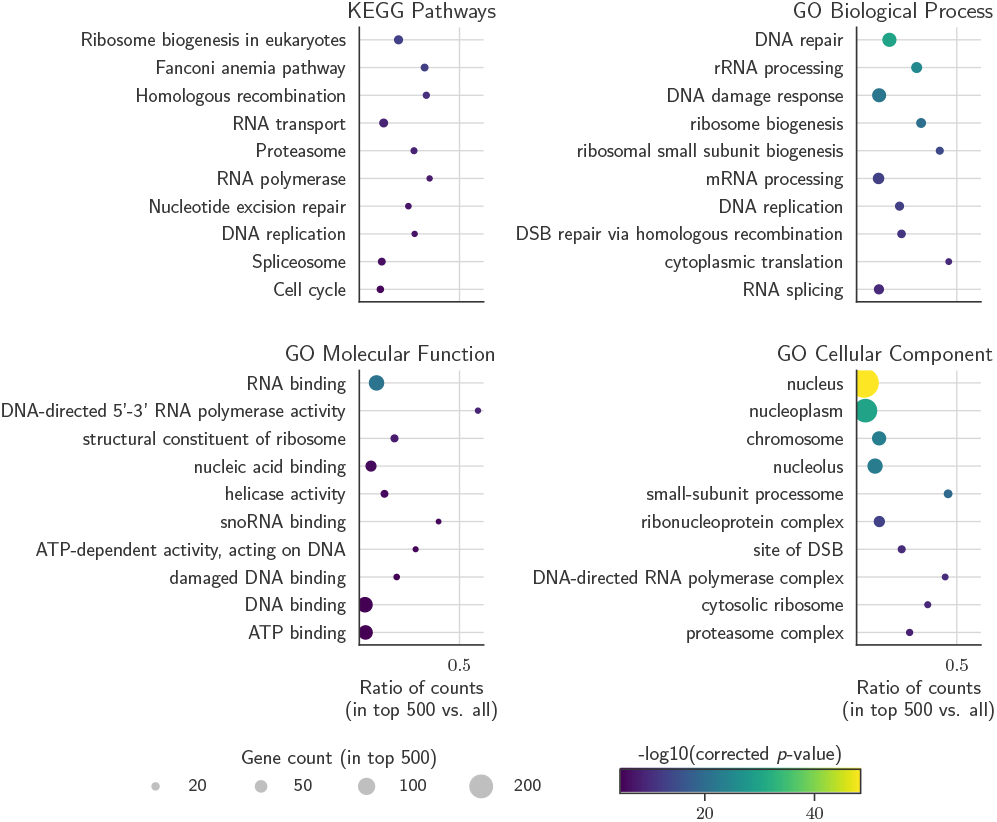
Top 10 enriched KEGG pathways and GO terms among the top 500 genes ranked by MUSICiAn across targets for the genome-wide MUSIC dataset. Top left to bottom right - KEGG pathways, GO biological processes, GO molecular functions, and GO cellular components. The horizontal axis shows the ratio between the numbers of genes annotated with the pathway or GO term among the top 500 ranked genes vs. all genes. Circle color denotes the negative log10 of the FDR-corrected *p*-value, and circle size indicates the number of genes annotated with a pathway or GO term among the top 500 ranked genes.

Functional enrichment analysis of the top 500 genes using GO annotations revealed links with repair-related biological processes (Fig. 5), including “DNA repair”, “double-strand break repair”, “double-strand break repair via homologous recombination”, and “interstrand cross-link repair”, further reinforcing the ability of MUSICiAn scores to capture and prioritize effects of genes on mutational spectra following the repair of CRISPR-induced DSB sites. Regarding molecular function, various binding activities, including DNA, damaged DNA, and ubiquitin-like protein ligase binding, as well as single-strand DNA helicase activity were identified, all functions required for DNA damage signalling and repair (Zhao *et al*., 2020; Schwertman et al., 2016; Pederiva et al., 2016) (Fig. 5).

Overall, MUSICiAn recovered known patterns and associations relevant to the repair of double-strand DNA breaks. While the AP performance may appear modest, it is significantly better than random. Nevertheless, mutational spectra exhibited relatively low coverage per sgRNA (median: MUSIC 2361.08 vs. Repair-seq 565201.97), leading to noisier mutational spectra that posed additional challenges in differentiating between true repair factors and noisy samples. Moreover, the assumption that mutational spectra deviating from the expected wild-type arise upon silencing of genes associated with DNA repair does not preclude the existence of other genes involved in DNA repair that do not affect mutational spectra. Such genes may not play a central role in the pathway, or their loss of function may be compensated by other genes, resulting in smaller effects and appropriately lower MUSICiAn rankings, while negatively biasing the AP.

### MUSICiAn identifies lesser-appreciated players in DSB repair

After analyzing established genes and pathways, we also examined several lesser-recognized pathways and processes emerging from the MUSICiAn analysis of the MUSIC dataset. Intriguingly, “Ribosome biogenesis in eukaryotes” was the top enriched KEGG pathway (Fig. 5), aligning with emerging literature from the last decade suggesting a potential cross-talk between ribosome biogenesis and DNA repair pathways (Ogawa and Baserga, 2017). Recent studies have also implicated the nucleolus, a major site of ribosome synthesis and the top enriched cellular component, in the regulation of cellular processes, including DNA repair (Lindström *et al*., 2018; Scott and Oeffinger, 2016; Korsholm *et al*., 2020).

The proteasome and spliceosome were additionally identified as enriched pathways. The proteasome plays a role in the regulation of the *Rnf8*-*Rnf168* pathway, which itself works to recruit repair factors to DSB sites (Schwertman *et al*., 2016; Krogan *et al*., 2004), and the inhibition of which has been previously shown to reduce HDR events (Cron *et al*., 2013). As for the spliceosome, there is growing evidence of a role in DNA repair, with studies suggesting that splicing regulates the expression of *Rnf8*, further controlling ubiquitin-signaling at DSBs (Pederiva *et al*., 2016).

### Enriched pathways promote homology-directed repair

We analyzed how the genes in the identified pathways influenced the frequencies of different mutation types by fitting a linear regression model per pathway, mutation type, and target site, and using the mutation type frequency per gene knockout and target site as response variable. Some pathways lacked sufficient gene representation to fit a reliable regression model (*<* 3 samples) and were excluded on a per-analysis basis (Fig. 6).

**Fig. 6:**
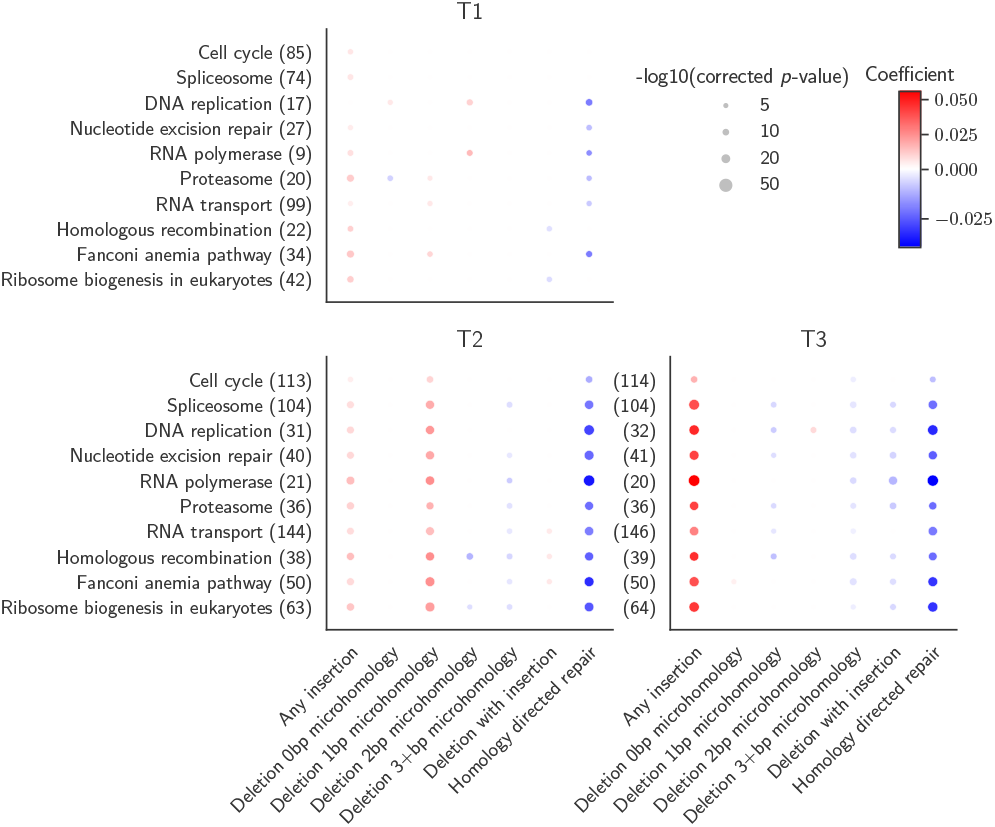
Effect of genes annotated with the top 10 enriched pathways on mutation frequency. We considered all genes in the top 10 pathways enriched amongst the top 500 genes ranked by MUSICiAn based on the genome-wide MUSIC spectra, with the final number of genes available per target dependent on the quality of the obtained mutational spectra. Each dot denotes a linear regression analysis of gene effect on mutation frequency per term or pathway (vertical axis, with gene count), mutation category (horizontal axis), and target site (panels for targets T1-T3 in MUSIC). Dot color denotes the regression coefficient, and dot size indicates the negative log10 of the FDR-corrected *p*-value. Points with non-significant corrected *p*-values (*>* 0.05) were excluded.

Based on the fitted models, we observed that the genes in each of the enriched pathways promoted HDR events and repressed insertion events across the target sites in the MUSIC genome-wide screen (Fig. 6, T1, T2, T3). Since NHEJ has been associated with introducing insertions at CRISPR-induced DSB sites (Lieber, 2008; Molla and Yang, 2020), we suggest that the rise and fall in the frequency of insertion and HDR events could reflect a change in the fraction of DSBs repaired via the NHEJ and HDR pathways. On the other hand, patterns pertaining to the promotion or inhibition of deletion events with or without MH were more sequence-context dependent, making it difficult to associate an inhibited pathway with how it might influence NHEJ and MMEJ. We note that the additional variation exhibited by MUSIC target site T1 could be an artifact of the noisier mutational profiles obtained for that target.

### MUSICiAn identifies novel gene-DSB repair associations

Analysis of the top 5 genes ranked by MUSICiAn for the genome-wide MUSIC dataset (Fig. 7) revealed two well-known DSB repair genes, *H2ax* (Scully and Xie, 2013) and *Xrcc5* (Taccioli *et al*., 1994). The others three genes, *Atp6v1g1, Metap2*, and *H2ac18*, were not annotated with the “double-strand break repair” GO term. The top-ranked gene was *Atp6v1g1*, for which one other study has reported an effect on HDR repair frequency after knockdown of *Atp6v1g1* via RNA interference (Adamson *et al*., 2012). The MUSIC spectra for target sites T2 and T3 showed a relative decrease in the frequency of HDR events after CRISPR knockout of *Atp6v1g1* compared to the geometric mean of the pseudo-controls. A similar tendency was observed for *Metap2*, a gene associated with ribosomal activity, and for *H2ac18*, a histone gene. Identifying histones is not surprising, as the chromatin state regulates DNA damage response by modulating accessibility to DNA damage sites by repair factors (Hunt *et al*., 2013). However, to our knowledge, no previous studies have identified an influence of *Metap2* or *H2ac18* on DNA repair pathways or HDR in particular. Further experimentation will be required to validate the impact of these top-ranking genes on mutational spectra and to investigate their role within the DSB repair process.

**Fig. 7:**
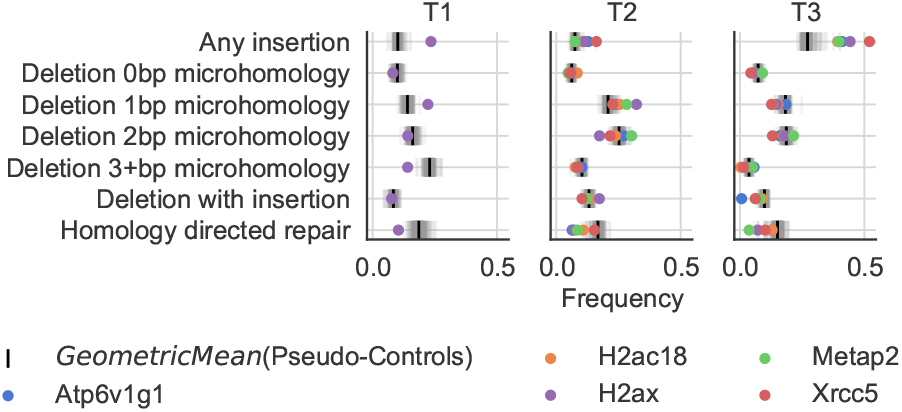
Mutational spectra of the top 5 genes ranked by MUSICiAn based on the genome-wide MUSIC screen. Colored dots denote the frequency obtained under knockout of the indicated gene The vertical axis shows mutation types. The horizontal axis shows frequency. The grey lines represent 300 randomly sampled genes. The black lines show the geometric mean of the pseudo-controls. The colored dots show the top genes. Some dots are not shown for T1, as the sgRNAs were filtered out during quality analysis.

## Conclusion

In this work, we introduced MUSICiAn, a control-free method to identify genes involved in DSB repair from gene perturbation screens with mutational spectra readout. MUSICiAn is developed for genome-wide perturbation screens, and leverages the fact that most genes have negligible influence on DSB repair and mutational spectra to frame the discovery as an outlier detection task. The goal of MUSICiAn is to both estimate the central tendency and identify genes with outlying spectra by analyzing the distribution of all mutational spectra.

Pseudo-controls estimated by MUSICiAn provided a good approximation of the actual non-targeting controls available for the Repair-seq dataset, showing that MUSICiAn could also be effective at sub-genome scale, provided the assumption that most genes have minimal effect on the spectra can reasonably be made. Notably, the combination of ILR transformation and robust covariance used by MUSICiAn contributed to an improved estimation of the central tendency and pseudo-controls.

Further MUSICiAn analysis of the genome-wide MUSIC data demonstrated an ability to recover known DSB repair genes and suggest candidates for further investigation, including *Atp6v1g1, Metap2*, and *H2ac18*. Our findings indicated that genes involved in ribosome biogenesis, the proteasome, and the spliceosome could play a significant role in modulating the frequency of HDR events, suggesting their involvement in DSB repair.

Obtaining sufficient coverage in genome-wide perturbation studies with sequence-based output remains a challenge that has also been noted in prior studies (Hussmann *et al*., 2021). Low coverage could limit the ability to detect subtle changes in mutagenic activity for rarer outcomes as the data becomes too sparse. To address this, we chose to aggregate mutational outcomes into broader categories. However, MUSICiAn could be applied with any collection of outcomes, as fine-grained as desired, and as the resolution across the different outcomes allows.

Overall, the results of MUSICiAn on the Repair-seq and the genome-wide MUSIC datasets highlighted that the method can effectively estimate pseudo-controls and identify genes with an impact on mutational spectra, enabling analyses of large-scale screens where designing realistic controls may be challenging.

## Acknowledgements

The authors acknowledge the High-Performance Compute cluster of the INSY department at Delft University of Technology. The illustrations in this article were created using BioRender.com.

## References

Adamson, B., et al. (2012). A genome-wide homologous recombination screen identifies the RNA-binding protein RBMX as a component of the DNA-damage response. Nature cell biology, 14(3), 318–328.

Aggarwal, C. C. (2016). Outlier Analysis Second Edition.

Aitchison, J. (1983). Principal component analysis of compositional data. Biometrika, 70(1), 57–65.

Ashburner, M., et al. (2000). Gene Ontology: tool for the unification of biology. Nature genetics, 25(1), 25–29.

Awwad, S. W., et al. (2023). Revolutionizing dna repair research and cancer therapy with crispr–cas screens. Nature Reviews Molecular Cell Biology, pages 1–18.

Benjamini, Y. and Hochberg, Y. (1995). Controlling the False Discovery Rate: A Practical and Powerful Approach to Multiple Testing. Journal of the Royal statistical society: series B (Methodological), 57(1), 289–300.

Bothmer, A., et al. (2017). Characterization of the interplay between dna repair and crispr/cas9-induced dna lesions at an endogenous locus. Nature communications, 8(1), 13905.

Burgers, P. M. (1998). Eukaryotic DNA polymerases in DNA replication and DNA repair. Chromosoma, 107(4), 218–227.

Ceccaldi, R., et al. (2016). Repair pathway choices and consequences at the double-strand break. Trends in cell biology, 26(1), 52–64.

Chen, A. (2011). Parp inhibitors: its role in treatment of cancer. Chinese journal of cancer, 30(7), 463.

Clay, D. E. and Fox, D. T. (2021). DNA damage responses during the cell cycle: insights from model organisms and beyond. Genes, 12(12), 1882.

Consortium, T. G. O. (2021). The Gene Ontology resource: enriching a GOld mine. Nucleic acids research, 49(D1), D325–D334.

Cortez, D. (2019). Replication-coupled DNA repair. Molecular cell, 74(5), 866–876.

Cron, K. R., et al. (2013). Proteasome inhibitors block DNA repair and radiosensitize non-small cell lung cancer. PloS one, 8(9), e73710.

Egozcue, J. J., et al. (2003). Isometric Logratio Transformations for Compositional Data Analysis. Mathematical geology, 35(3), 279– 300.

Filzmoser, P., et al. (2009). Principal component analysis for compositional data with outliers. Environmetrics: The Official Journal of the International Environmetrics Society, 20(6), 621– 632.

Hunt, C. R., et al. (2013). Histone modifications and DNA double-strand break repair after exposure to ionizing radiations. Radiation research, 179(4), 383–392.

Hurov, K. E., et al. (2010). A genetic screen identifies the triple t complex required for dna damage signaling and atm and atr stability. Genes & development, 24(17), 1939–1950.

Hussmann, J. A., et al. (2021). Mapping the genetic landscape of DNA double-strand break repair. Cell, 184(22), 5653–5669.

Jenkins, T., et al. (2021). The HelQ human DNA repair helicase utilizes a PWI-like domain for DNA loading through interaction with RPA, triggering DNA unwinding by the HelQ helicase core. NAR cancer, 3(1), zcaa043.

Kanehisa, M., et al. (2023). KEGG for taxonomy-based analysis of pathways and genomes. Nucleic Acids Research, 51(D1), D587– D592.

Korsholm, L. M., et al. (2020). Recent advances in the nucleolar responses to DNA double-strand breaks. Nucleic acids research, 48(17), 9449–9461.

Krogan, N. J., et al. (2004). Proteasome involvement in the repair of DNA double-strand breaks. Molecular cell, 16(6), 1027–1034.

Lieber, M. R. (2008). The mechanism of human nonhomologous DNA end joining. Journal of Biological Chemistry, 283(1), 1–5.

Lindström, M. S., et al. (2018). Nucleolus as an emerging hub in maintenance of genome stability and cancer pathogenesis. Oncogene, 37(18), 2351–2366.

Lubbe, S., et al. (2021). Comparison of zero replacement strategies for compositional data with large numbers of zeros. Chemometrics and Intelligent Laboratory Systems, 210, 104248.

Mahalanobis, P. C. (2018). On the generalized distance in statistics. Sankhya: The Indian Journal of Statistics, Series A (2008-), 80, S1–S7.

Michl, J., et al. (2016). Interplay between fanconi anemia and homologous recombination pathways in genome integrity. The EMBO journal, 35(9), 909–923.

Molla, K. A. and Yang, Y. (2020). Predicting CRISPR/Cas9-induced mutations for precise genome editing. Trends in biotechnology, 38(2), 136–141.

O’Connell, B. C., et al. (2010). A genome-wide camptothecin sensitivity screen identifies a mammalian mms22l-nfkbil2 complex required for genomic stability. Molecular cell, 40(4), 645–657.

Ogawa, L. and Baserga, S. (2017). Crosstalk between the nucleolus and the dna damage response. Molecular bioSystems, 13(3), 443–455.

Olivieri, M., et al. (2020). A genetic map of the response to dna damage in human cells. Cell, 182(2), 481–496.

Pederiva, C., et al. (2016). Splicing controls the ubiquitin response during DNA double-strand break repair. Cell Death & Differentiation, 23(10), 1648–1657.

Schwertman, P., et al. (2016). Regulation of DNA double-strand break repair by ubiquitin and ubiquitin-like modifiers. Nature reviews Molecular cell biology, 17(6), 379–394.

Scott, D. D. and Oeffinger, M. (2016). Nucleolin and nucleophosmin: nucleolar proteins with multiple functions in dna repair. Biochemistry and cell biology, 94(5), 419–432.

Scully, R. and Xie, A. (2013). Double strand break repair functions of histone H2AX. Mutation Research/Fundamental and Molecular Mechanisms of Mutagenesis, 750(1-2), 5–14.

Scully, R., et al. (2019). Dna double-strand break repair-pathway choice in somatic mammalian cells. Nature reviews Molecular cell biology, 20(11), 698–714.

Shen, M. W., et al. (2018). Predictable and precise template-free crispr editing of pathogenic variants. Nature, 563(7733), 646–651.

Shou, J., et al. (2018). Precise and predictable crispr chromosomal rearrangements reveal principles of cas9-mediated nucleotide insertion. Molecular cell, 71(4), 498–509.

Smogorzewska, A., et al. (2010). A genetic screen identifies fan1, a fanconi anemia-associated nuclease necessary for dna interstrand crosslink repair. Molecular cell, 39(1), 36–47.

Taccioli, G. E., et al. (1994). Ku80: product of the XRCC5 gene and its role in DNA repair and V (D) J recombination. Science, pages 1442–1445.

Thomas, A., et al. (2022). Division of Labor by the HELQ, BLM, and FANCM Helicases during Homologous Recombination Repair in Drosophila melanogaster. Genes, 13(3), 474.

Trenner, A. and Sartori, A. A. (2019). Harnessing dna double-strand break repair for cancer treatment. Frontiers in oncology, 9, 1388.

van Overbeek, M., et al. (2016). Dna repair profiling reveals nonrandom outcomes at cas9-mediated breaks. Molecular cell, 63(4), 633–646.

van Schendel, R., et al. (2022). SIQ: easy quantitative measurement of mutation profiles in sequencing data. NAR Genomics and Bioinformatics, 4(3), qac063.

Ward Jr, J. H. (1963). Hierarchical Grouping to Optimize an Objective Function. Journal of the American statistical association, 58(301), 236–244.

Wyatt, D. W., et al. (2016). Essential roles for polymerase θ-mediated end joining in the repair of chromosome breaks. Molecular cell, 63(4), 662–673.

Zhang, Y., et al. (2009). Involvement of nucleotide excision and mismatch repair mechanisms in double-strand break repair. Current genomics, 10(4), 250–258.

Zhao, F., et al. (2020). DNA end resection and its role in DNA replication and DSB repair choice in mammalian cells. Experimental & Molecular Medicine, 52(10), 1705–1714.

Zimmermann, M., et al. (2018). Crispr screens identify genomic ribonucleotides as a source of parp-trapping lesions. Nature, 559(7713), 285–289.

