## Supplementary Materials for "MUSICiAn: Genome-wide Identification of Genes Involved in DNA Repair via Control-Free Mutational Spectra Analysis"

#### Contents

**1 Supplementary Tables** **2**

**2 Supplementary Figures** **4**

### 1 Supplementary Tables

| Target | Sample | **Before** | Step *i* | Step *ii* | Step *iii* | **Step *iv*** |
|---|---|---|---|---|---|---|
| T1 | MB01 | **89414** | 75218 | 74654 | 64700 | **60975** |
| | MB02 | **89423** | 76352 | 75789 | 67200 | **63347** |
| T2 | MB03 | **89492** | 80058 | 79887 | 79606 | **79401** |
| | MB04 | **89481** | 82096 | 81926 | 80934 | **80646** |
| T3 | MB05 | **89477** | 79296 | 79042 | 78796 | **78675** |
| | MB06 | **89478** | 78492 | 78237 | 78112 | **78037** |

Table S1: Breakdown of sgRNA counts after each QA filtering step as described in "Quality analysis and sgRNA filtering" from the main article. Briefly, we filtered out sgRNAs per replicate by applying the following criteria in order: *(i)* a total mutated read count below 700; *(ii)* only two sgRNA representations for the gene; *(iii)* a median pairwise Pearson correlation coefficient below 0.6 compared to other sgRNAs for the same gene within the same replicate; or *(iv)* a median pairwise Pearson correlation below 0.6 compared to paired replicates at the same target site.

| Target | Sample | **Before** | Step *i* | Step *ii* | Step *iii* | **Step *iv*** |
|---|---|---|---|---|---|---|
| T1 | MB01 | **18406** | 18023 | 17887 | 17130 | **17004** |
| | MB02 | **18405** | 18025 | 17889 | 17251 | **17135** |
| T2 | MB03 | **18406** | 18167 | 18116 | 18090 | **18089** |
| | MB04 | **18406** | 18264 | 18213 | 18144 | **18140** |
| T3 | MB05 | **18406** | 18224 | 18149 | 18134 | **18133** |
| | MB06 | **18406** | 18178 | 18103 | 18095 | **18092** |

Table S2: Breakdown of gene counts after each QA filtering step per the steps outlined in Table S1 above.

| **MUSICiAn Category** | **SIQ Categories** | **Description** |
|---|---|---|
| Wild-type | WT | No mutation. |
| Deletion with insertion | DELINS<br>TINS<br><br><br>TANDEMDUPLICATION<br><br><br>TANDEMDUPLICATION-_COMPOUND | Deletion with insertion.<br>Deletion with an insertion where the insert is copied from the flank.<br>Duplication of sequence immediately flanking the cut-site.<br>A tandem duplication with some additional inserted sequence. |
| Insertion | INSERTION | Any simple insertion event. |
| Homology-directed repair | HDR | Homology-directed repair event. |
| Deletion with no microhomology<br>Deletion with 1bp microhomology<br>Deletion with 2bp microhomology<br>Deletion with 3+bp microhomology | DELETION | Any simple deletion event. Additional details recorded include the presence and length of any homology between one side of the deleted sequence and the opposing flank of the cut site. |

Table S3: Mapping of MUSICiAn mutation categories to SIQ mutation types and details.

#### 2 Supplementary Figures

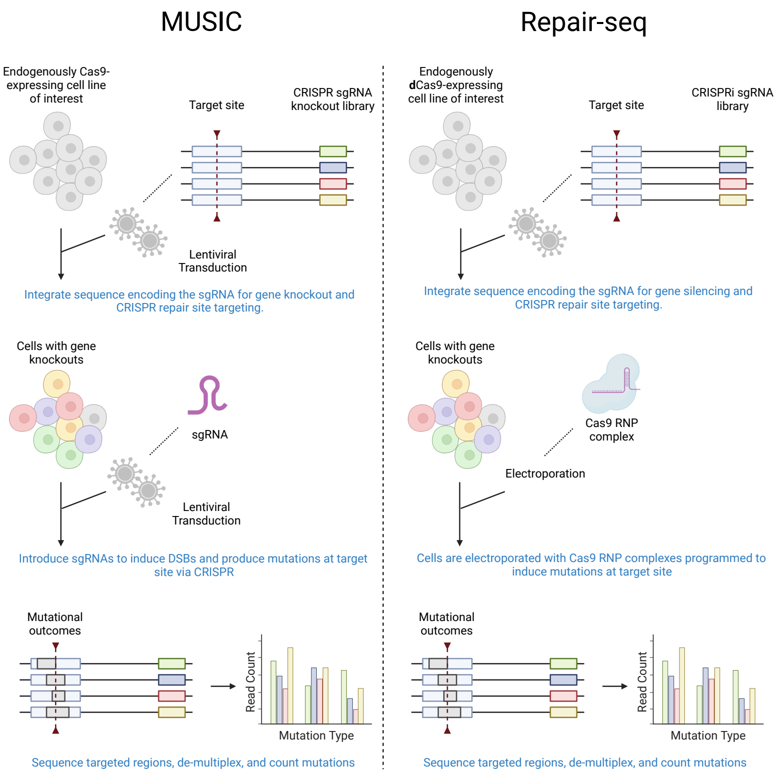


Figure S1: Illustration of CRISPR gene perturbation screens with mutational spectra readout. First, sequences are integrated into the genomes of cells via lentiviral transduction. Each sequence contains two elements: *(i)* a sgRNA-encoding region to knockout (MUSIC) or silence (Repair-seq) a single gene, and *(ii)* a region common to all integrated sequences to be targeted with CRISPR to produce the mutational spectra. After genomic integration, several days of cell culture are allowed for genes to be knocked out. Following this, MUSIC again uses lentiviral transduction to introduce sgRNAs targeting the common region to the Cas9-expressing cells. Repair-seq uses electroporation to introduce Cas9 RNP complexes to the cells to induce DSBs at the target site. After allowing time for cell culture for DNA cleavage and repair, DNA sequencing was performed to capture the final CRISPR repair products.

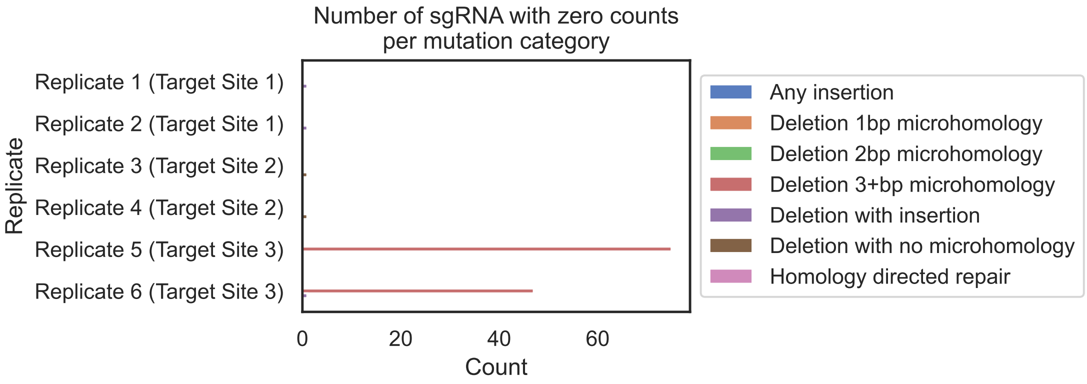


Figure S2: Counts of sgRNAs with zero values per mutation category and replicate.
